# Different Sensitivity to Ethanol and Sucrose in DAT and SERT Knockout Rats

**DOI:** 10.1101/2025.08.16.670641

**Authors:** Emile E. Zweistra, Anna Cabiscol-Claveria, Michel M.M. Verheij, Edine Remmers, Rick Hesen, Thomas A. Scholtes, Debbie R.M. Tesselaar, Arnt F.A. Schellekens, Jan Booij, Judith R. Homberg, Cyprien G. J. Guerrin

**Affiliations:** Department of Medical Neuroscience, Donders Institute for Brain, Cognition, and Behavior, Radboud University Medical Center, Nijmegen, the Netherlands; Department of Psychiatry, Radboud University Medical Center, Nijmegen, The Netherlands; Nijmegen Institute for Scientist-Practitioners in Addiction (NISPA), Nijmegen, The Netherlands; Department of Radiology and Nuclear Medicine, Amsterdam University Medical Centers, location Academic Medical Center, Amsterdam, The Netherlands; Department of Medical Imaging, Radboud University Medical Center, Nijmegen, The Netherlands; Experimental Pharmacopsychology and Psychological Addiction Research, Department of Adult Psychiatry and Psychotherapy, University Hospital of Psychiatry Zurich, University of Zurich, Zurich, Switzerland

**Author notes:** Corresponding author: Cyprien G. J. Guerrin, Department of Medical Neuroscience, Donders Institute for Brain, Cognition, and Behavior, Radboud University Medical Center, Nijmegen, the Netherlands. Authors contributed equally (Emile E. Zweistra, Anna Cabiscol-Claveria). Authors jointly supervised this work (Judith R. Homberg, Cyprien G. J. Guerrin).

## Abstract

**Background:** Dopamine and serotonin are key regulators of reward sensitivity, yet their distinct roles in motivating natural (e.g., sucrose) versus drug (e.g., ethanol) rewards remain unclear. Understanding these mechanisms could help explain individual variability in reward processing relevant to substance use vulnerability.

**Methods:** We assessed reward sensitivity in dopamine transporter (DAT) and serotonin transporter (SERT) knockout (KO) rats using both home cage (two-bottle choice for sucrose and ethanol) and operant paradigms (Pavlovian and instrumental learning).

**Results:** DAT KO rats showed lower sucrose preference (-27% for 2%, -13% for 4%) and intake (-42% for 4%), diminished Pavlovian responding for sucrose (-68%), and slower acquisition of sucrose-taking behavior (∼+30 days) compared to WT rats. DAT KO rats also showed reduced ethanol preference in the home cage (-16%) despite an unchanged intake. Furthermore, operant performed was markedly reduced operant performance after the sucrose-to-ethanol transition (-83%), with no increase in ethanol-taking following sucrose exposure (0% change), unlike WT controls (+41%).

SERT KO rats presented reduced sucrose preference (-5%) and intake (-46%) for the 4% solution only. In addition, SERT KO rats also showed reduced Pavlovian sucrose responding (- 28%) and slower acquisition of sucrose-taking (∼+30 days) but intact responding and learning for ethanol. In the home cage, they displayed lower ethanol preference (-35%) without significant change in operant ethanol performance. A modest overall increase in ethanol-taking was seen post-sucrose in both SERT KO and WT, but without genotype-specific effects.

**Conclusion and Implications:** DAT deletion broadly impaired sensitivity for both natural sucrose and ethanol rewards, particularly under effortful or devalued conditions. In contrast, SERT deletion produced more selective impairments by disrupting sucrose operant responding and moderately reducing ethanol reward preference. These findings reveal distinct but overlapping roles of DAT and SERT in regulating reward sensitivity, with implications for understanding individual vulnerability to substance use.

## 1. Introduction

The capacity to experience pleasure and motivation from rewarding stimuli i.e. reward sensitivity, influences learning, decision-making, motivation and emotion functioning (Pulver et al., 2020). Aberrant reward sensitivity, whether excessive or blunted, is involved in the pathophysiology of addiction, major depression, and anxiety (Bart et al., 2021; Berry et al., 2019; Pulver et al., 2020; Tatnell et al., 2019; Volkow et al., 2010). Longitudinal studies demonstrate that individuals who report stronger acute euphoria and stimulation from alcohol are more likely to develop alcohol-use disorder (AUD) later in life (King et al., 2016, 2014, 2011; Morean and Corbin, 2010; Parker et al., 2020). Sweet preference follows a similar risk pattern: persons with a family history of AUD find sweet tastes more pleasant and display greater impulsivity and reward-related brain activation (Carroll et al., 2002; Han et al., 2020; Kampov-Polevoy et al., 2004, 2001, 1999; Weafer et al., 2014). Conversely, diminished reward sensitivity contributes to the anhedonia and motivational deficits characteristic of substance use disorders and depression (Hausman et al., 2018). Reward sensitivity is therefore a multidimensional trait that varies widely both within and between diagnostic conditions.

Central to reward sensitivity are the dopaminergic and serotonergic neurotransmitter systems (Berger et al., 2009; Juárez Olguín et al., 2016). Dopamine release enhances motivation and reward sensitivity, thereby reinforcing behavior like sweet food and alcohol consumption (Bari et al., 2010; Deyoung, 2013; Mkrtchian et al., 2025), while serotonin release modulates reward anticipation, and punishment sensitivity (Bari et al., 2010; Fletcher et al., 1995; Homberg et al., 2014; Mkrtchian et al., 2025). Dysregulation of these systems is associated with various psychiatric disorders such as AUD, major depression, anxiety disorders, and attention-deficit/hyperactivity disorder (Cabana-Domínguez et al., 2022; Franco et al., 2021; Klein et al., 2019; Lin et al., 2014; Pourhamzeh et al., 2022; Speranza et al., 2025). The dopamine transporter (DAT) and serotonin transporter (SERT) play an important role in the reuptake of synaptic dopamine and serotonin, respectively, positioning them as critical targets for pharmacological intervention. Yet, their precise contributions in modulating reward sensitivity such as sweet tastes and alcohol remain unclear. Clarifying the roles of DAT and SERT in these processes may offer valuable insights into novel therapeutic strategies for disorders involving altered reward sensitivity.

Genetically modified animals, such as DAT and SERT knockout (KO) rats, offer valuable tools to investigate these mechanisms directly. Reward sensitivity in rodents can be assessed robustly using paradigms involving rewards with distinct properties, such as sucrose (sweet solution) and ethanol (alcohol). DAT KO rats exhibit chronically elevated extracellular dopamine levels (Leo et al., 2018), yet paradoxically show reduced sensitivity to various natural rewards, including social interaction, cheese preference and sucrose preference in the sucrose preference test (SPT) (Cinque et al., 2018; Mallien et al., 2022). However, findings across species and conditions are mixed as DAT KO mice exhibit increased sucrose preference and consumption in the SPT (Perona et al., 2008), while DAT knockdown mice show reduced ethanol intake and preference (24-hour home cage drinking) without changes in saccharin intake (Bahi and Dreyer, 2019). Similarly, SERT KO rodents, which have elevated extracellular serotonin, exhibit mixed responses: some studies report no or only minor reductions in sucrose preference and intake in SERT KO males (Bearer et al., 2009; Diniz et al., 2021; Perona et al., 2008; Sun et al., 2024), while others find reduced intake, with preference reduction observed only in females (Olivier et al., 2008). Operant studies also show no difference in progressive ratio responding, but reduced sucrose intake with repeated long-access (6h) (Karel et al., 2020). Notably, SERT KO mice show reduced ethanol motivation with lower breakpoints in progressive ratio schedules and increasing fixed ratio schedule, as well as reduced ethanol preference and intake in two-bottle choice tests, despite similar post-prandial ethanol consumption (Bearer et al., 2009; Hussain et al., 2022; Kelaï et al., 2003; Lamb and Daws, 2013).

These discrepancies highlight significant gaps in understanding, especially regarding the comparative motivational and rewarding values of ethanol and sucrose. Crucially, no prior studies have directly compared ethanol and sucrose reward processing in the same DAT and SERT KO models using controlled operant paradigms. Addressing these knowledge gaps is essential to precisely characterize reward-specific alterations associated with DAT and SERT dysfunction, thereby advancing our comprehension of underlying neural mechanisms and informing targeted therapeutic interventions.

This study addresses these knowledge gaps by directly comparing sucrose and ethanol reward sensitivity in DAT and SERT KO rats and their wild type (WT) counterparts. Sucrose and ethanol preference was assessed in the home cage, and motivation was quantified using operant paradigms for each reward type separately. To our knowledge, this is the first study to systematically compare operant responding for ethanol and sucrose within the same genetically modified animal models, offering novel insights into reward-type-specific deficits and the translational relevance of transporter dysfunction.

## 2. Materials and Methods

### 2.1. Animals

All experimental procedures were conducted in compliance with the European Directive 2010/63/EU and the Law on Animal Experiments in the Netherlands and were approved by the Centrale Commissie Dierproeven (CCD; permit number: AVD10300 2023 16779). A total of 36 male rats were included in the study, divided into four experimental groups: 10 DAT WT, 10 DAT KO, 8 SERT WT, and 8 SERT KO. The SERT and DAT KO rats were generated on a Wistar Unilever and Wistar-Han genetic background, respectively, as previously described (Homberg et al., 2007; Leo et al., 2018), and all animals were bred and housed in the animal facility of Radboud University (Nijmegen, Netherlands). Rats were group-housed in standard type IV cages, with 2-3 animals per cage, under a reverse 12:12 light-dark cycle (lights on at 07:30 pm) with food and water provided ad libitum. Animals were housed exclusively with cage-mates of the same genotype and genetic background (e.g., DAT WT with DAT WT, SERT KO with SERT KO) to prevent potential social stress or dominance hierarchies between different genotypes or strains. Each cage was enriched with bedding, gnaw sticks, and shelters. Housing conditions were maintained at a temperature of 21–22 °C ± 1 °C and a relative humidity of 45–65%. Body weight was monitored weekly.

### 2.2. Chemicals

A 15% ethanol solution was prepared by diluting 100% absolute ethanol (Biosolve B.V., Valkenswaard, NL) in autoclaved drinking water. This solution was used for all ethanol-related experiments and prepared once a week.

Sucrose solutions (2% and 4%) were prepared in autoclaved drinking water one day before testing. A 10% solution was prepared weekly for operant sessions.

### 2.3. Experimental design

The experimental design consisted of three main phases, as illustrated in Figure 1A. The first two parts consisted of a home-cage two-bottle choice paradigm to evaluate voluntary consumption of a sucrose as well as ethanol solution versus water, consisting of two 6-hour sessions for sucrose exposure and fourteen 24-hour sessions for ethanol exposure across a 7-week period. Finally, animals underwent operant conditioning over a 23–25 week period, which included both Pavlovian and instrumental (taking) phases for ethanol and sucrose rewards.

**Figure 1.**
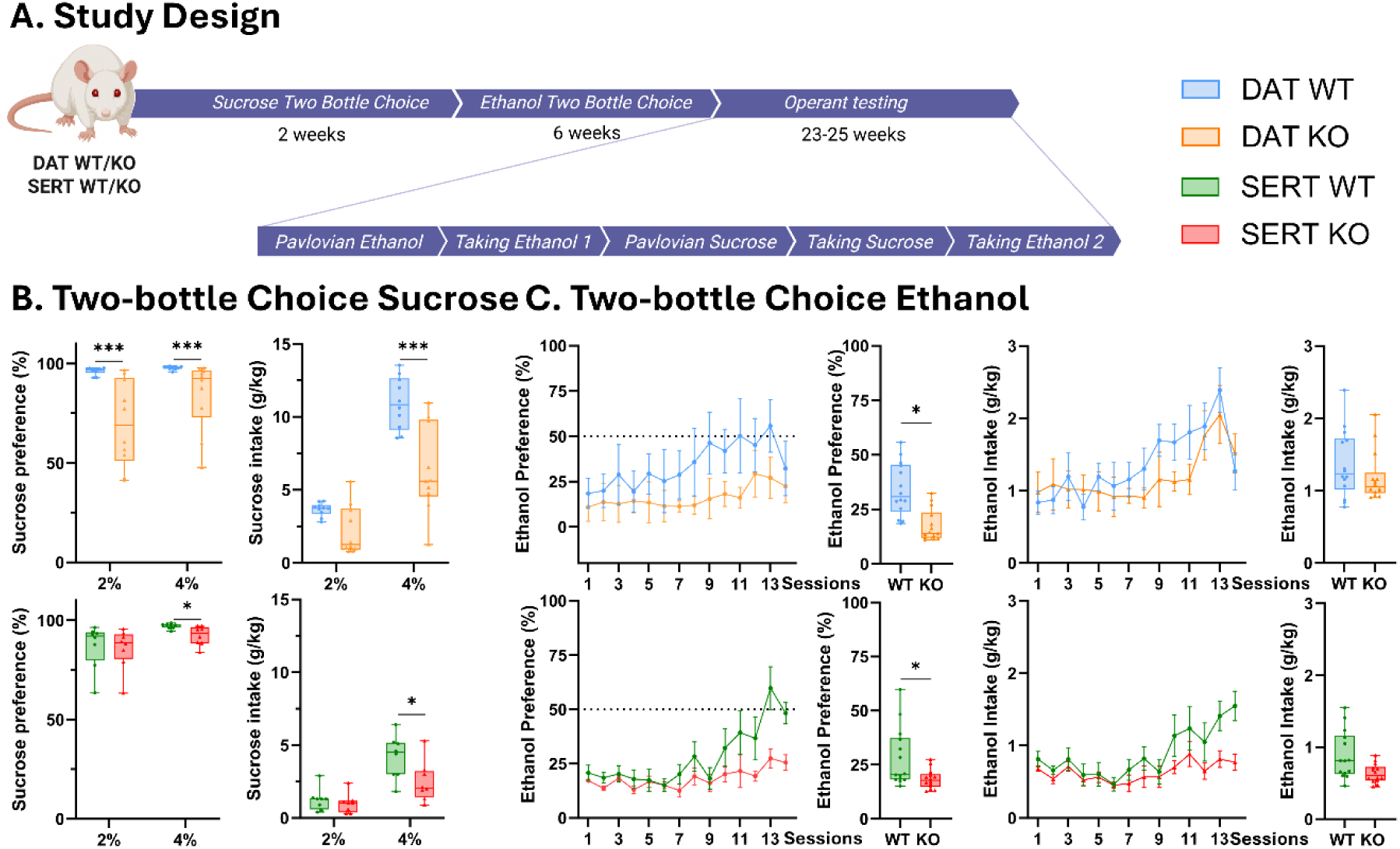
Study design and behavioral outcomes across DAT and SERT cohorts. (A) Experimental timeline illustrating the sequence of behavioral testing across DAT and SERT cohorts. (B) Sucrose preference and intake results for 2% (second session) and 4% sucrose solutions in the home cage. (C) 15 % ethanol preference and intake results across 14 sessions in the home cage. Data are presented as boxplots (minimum to maximum) or mean±SEM. Statistically significant differences between groups are indicated by asterisks: *p < 0.05, **p < 0.01, ***p < 0.001. WT = wild-type; KO = knockout. N = 10 DAT WT, 10 DAT KO, 8 SERT WT, 8 SERT KO.

### 2.4. Two-bottle choice sucrose consumption

Voluntary sucrose preference and intake were assessed using a two-bottle choice sucrose paradigm in the home cage before ethanol exposure. The test lasted five consecutive days. During the first two days, rats were habituated to four water bottles in their homecage. On days 3-5, rats were separated within their home cage using transparent dividers for ∼7.5 hours per day (during their active phase) and presented with one bottle of water and one bottle of sucrose solution (2%, 2%, and 4%, respectively). Bottle positions alternated per session to avoid bias formation. Fluid intake was measured by weighing the bottles before and after sessions. Sucrose preference was calculated as a percentage of sucrose intake relative to total fluid intake (Sucrose preference = (sucrose intake/sucrose intake + water intake) * 100). Sucrose intake was calculated per rat by multiplying the volume (mL) of sucrose solution consumed by its concentration (2% or 4%) and dividing the result by the rat’s body weight (in kg). Sucrose intake (g/kg) = (concentration [g/mL]*volume consumed [mL]/rat weight [kg]).

### 2.5. Two-bottle choice ethanol consumption

Voluntary ethanol preference and intake were assessed using an intermittent two-bottle choice paradigm in the home cage across 14 sessions, each lasting 24 hours. In each session, rats had simultaneous access to two bottles: one containing 15% ethanol and the other water. Bottle positions alternated each session to prevent side bias. Fluid consumption was calculated by weighing bottles before and after each session. Ethanol preference was calculated as the percentage of ethanol consumed relative to total fluid intake per cage. As rats were housed in pairs or trios per cage without dividers (to prevent the possible impact of repeated isolation), and grouped by genotype (KO with KO, WT with WT), ethanol intake per rat (in g/kg body weight) was calculated by multiplying the volume of 15% ethanol consumed (in mL) by its density-adjusted concentration (0.11835 g/mL, based on a 15% solution and ethanol’s density of 0.789g/mL), and dividing the result by the combined body weight of the cage-mates (in kg). Ethanol intake (g/kg) = (0.11835*volume consumed [mL])/ cage weight [kg]).

### 2.6. Operant Testing

Operant testing was conducted in 12 chambers (MedAssociates, St. Albans, VT, USA), each equipped with two retractable levers, cue lights positioned above each lever, a house light, and a central magazine with a liquid dipper (MedAssociates, St. Albans, VT, USA, ENV-202M-S). An infrared beam across the magazine recorded head entries (beam breaks), which served as a measure of consummatory behavior. Testing sessions were conducted on weekdays over a period of 23 weeks for the SERT cohort and 25 weeks for the DAT cohort. Before and after operant testing, rats had ad libitum access to food and water, ensuring that operant responses reflected hedonic and motivational factors rather than a compensation for food or water deprivation. The operant testing protocol consisted of five sequential phases: Pavlovian Conditioning Phase with Ethanol – Taking Phase with Ethanol (1st exposure) – Pavlovian Conditioning Phase with Sucrose – Taking Phase with Sucrose – Taking Phase with Ethanol (2nd exposure). Each phase is described in detail below.

#### Pavlovian Conditioning Phase - Ethanol and Sucrose

During this phase, a visual cue (cue light) was continuously illuminated above the retracted lever throughout the session. Each session began with a 60-second acclimatization period. Each session lasted 1 hour and consisted of 172 trials, during which 0.116 mL of 15% ethanol or 10% sucrose was delivered via a liquid dipper every twenty seconds without inter-trial intervals. A trial was considered successful (i.e., a completed trial) if the rat made a head entry into the magazine (beam break) while the reward was available. Rats progressed to the next phase after completing successfully at least 25 trials (completed trials) for two consecutive days. A minimum of five sessions were conducted for ethanol and three for sucrose. The number of completed trials was measured for each session.

#### Taking Phase - Ethanol and Sucrose

In this phase, rats were trained to perform an operant response under a fixed-ratio 1 (FR1) schedule to obtain either ethanol or sucrose. Each session lasted until the animal received 45 rewards or 1 hour elapsed, whichever occurred first. Sessions began with a 60-second acclimatization period. Each trial started with the insertion of a single lever (randomly assigned to the left or right side at the first session, counterbalanced across individuals to control for side preference), accompanied with a 1-second cue light above the lever, which remained extended for a maximum of 60 seconds or until a lever press occurred. A successful lever press triggered a 1-second cue light above the lever, followed by delivery of 0.116 mL of 15% ethanol or 10% sucrose solution via a liquid dipper. A 20-second time-out followed each reward delivery, during which the lever was retracted. A trial was considered completed when the rat pressed the lever and subsequently made a head entry into the magazine (beam break) while the reward was available. Rats progressed to the next phase or were removed from further testing in the final ethanol phase after achieving at least 25 completed trials on two consecutive days. The number of completed trials were recorded during each session.

### 2.7. Statistical Analysis

All behavioral data were analyzed using R version 4.5.0 (R Core Team, 2025). Normality and homogeneity of variance were assessed using visual inspection of residuals and formal tests (Shapiro–Wilk, Levene’s test, respectively). When assumptions were met, linear mixed effects models were used with WT as reference for the home cage two bottle choice, pavlovian phase and ethanol transition. To assess interaction effects (genotype*sessiontype) in the pavlovian phase, a likelihood ratio test (LRT) was performed by comparing the full model to a reduced model (without the interaction). Significant interactions were followed by post hoc comparisons using estimated marginal means with Bonferroni correction for multiple testing. For the sucrose two bottle choice, considering the change in task conditions (2% - 4%) and the violated homoscedasticity assumption, we used the non-parametric Mann-Witney U test (MWU) for between-group comparisons. For the independent taking phases in the operant chamber, a Kaplan–Meier survival analysis was conducted to assess the time it took animals to reach our predefined performance criteria: 25 completed trials over 2 consecutive days. Some animals did not reach the required criteria and therefore did not progress through all operant chamber phases. Data were missing for 5% of total sessions in DAT KO, 7% in DAT WT, 5% in SERT KO, and 8% in SERT WT rats. Two DAT KO animals were excluded from operant analyses due to humane endpoints being reached before the end of the study. All tests were two-tailed, with statistical significance set at p < 0.05. Detailed statistics results can be found in supplementary tables 1-8.

## 3. Results

### 3.1. DAT and SERT KO rats show reduced bodyweight compared to their respective WT controls

Linear mixed effect modelling using genotype as between subject variable and animal ID as random effect showed a main effect of genotype in the DAT cohort (β = –145.977, *SE* = 11.87, *t*(18.01) = 12.321, *p* = <.001). In the SERT cohort, analysis revealed a main effect of genotype (β = –52.856, SE = 11.78, t(14) = -4.486, p < .001).

### 3.2. DAT and SERT KO rats show reduced sucrose preference and intake in the sucrose two-bottle choice compared to WT controls

Voluntary sucrose preference and intake was assessed using a two-bottle choice paradigm prior to ethanol exposure (Fig. 1B).

In the DAT cohort, independent parallel MWU tests revealed that DAT KO rats exhibited significantly lower sucrose preference than WT controls for both 2% (-27%) and 4% sucrose solution (-13%) [2%: W = 95, p = < .001; 4%: W = 93, p = < .001]. Furthermore, MWU tests revealed that DAT KO sucrose intake was significantly lower than WT during the 4% session (- 42%, W = 89, *p* = .0.002) but not the 2% session (W = 78, *p* = .07).

In the SERT cohort, the same MWU analysis revealed that SERT KO rats showed lower sucrose preference for the 4% solution (-5%; W = 56, *p* < .05) than SERT WT animals, whereas no differences were found for the 2% solution (W = 37, *p* = .645; Fig. 1B). Regarding intake, MWU tests revealed that SERT KO rats consumed significantly less sucrose than WT rats in the 4% session (-46%; W = 56, *p* < 0.05), while no differences were found on the 2% session (W = 44, *p* = .235; Fig.1B).

### 3.3. DAT and SERT KO rats exhibit reduced ethanol preference but not intake in the two-bottle choice test compared to their respective WT controls

Voluntary ethanol preference and intake (g/kg body weight) were assessed using a two-bottle choice paradigm over 14 intermittent sessions (Fig. 1C).

For the DAT cohort, linear mixed effects modelling revealed a significant main effect of genotype on ethanol preference, with DAT KO rats showing reduced ethanol preference compared to WT animals (β = –16.53, *SE* = 6.37, *t*(8) = –2.60, *p* = .014). In contrast, there was no significant main effect of genotype on average ethanol consumption g/kg (β = –0.36, *SE* = 0.59, *t*(8) = –0.61, *p* = .561).

In the SERT cohort, the analysis also revealed a significant main effect of genotype on ethanol preference, with SERT KO rats exhibiting lower preference compared to SERT WT controls (-35%, *b* = -9.78, *SE* = 3.89, *t*(6) = -2.51, *p* < .05; Fig. 1C), but no significant difference in average ethanol intake between genotypes (β = -0.51, *SE* = 0.31, *t*(6) = -1.64, *p* = .153; Fig. 1E).

### 3.4. DAT and SERT KO rats show reduced Pavlovian responding for sucrose but not for ethanol

Pavlovian conditioning sessions assessed voluntary consumption of ethanol and sucrose (Fig. 2B), using the number of completed trials as an index for reward-driven engagement.

**Figure 2.**
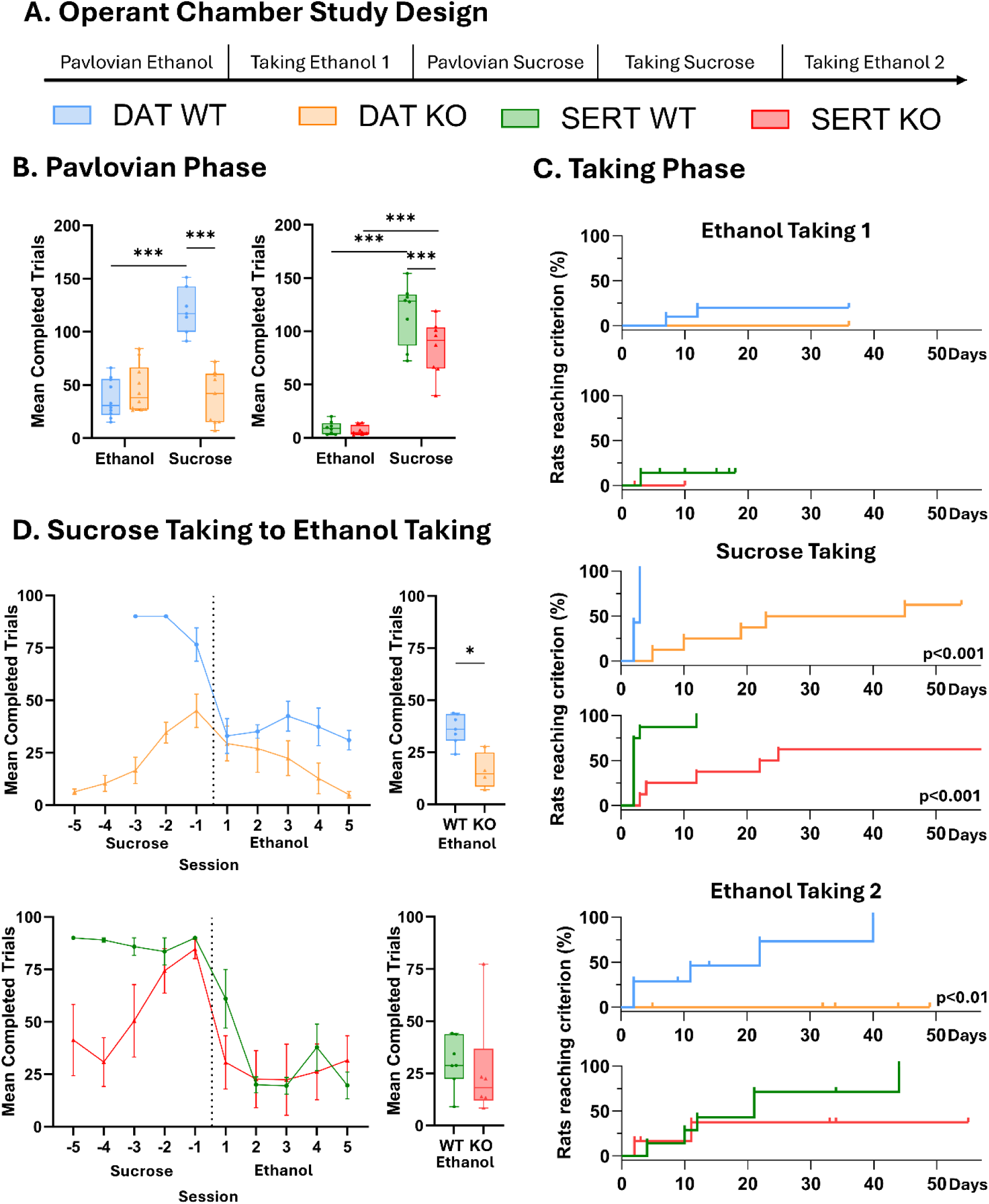
Operant chamber testing performance. (A) Timeline of operant chamber study design, indicating the order of Pavlovian and taking phases for ethanol and sucrose. (B) Mean completed trials during the Pavlovian phase for sucrose and ethanol conditioning, per 60-minute session, averaged across the same three sessions. N = 7-10 DAT WT, 9-10 DAT KO, 8 SERT WT, 8 SERT KO. (C) Days to reach criterion during each operant taking phase for ethanol and sucrose. N = 7 DAT WT, 5-8 DAT KO, 7-8 SERT WT, 4-8 SERT KO. (D) Completed trials during transition from sucrose taking to ethanol taking (±5 sessions), including boxplots of trial counts. N = 7 DAT WT, 5 DAT KO, 7 SERT WT, 6 SERT KO. Data are presented as boxplots (minimum to maximum) or mean±SEM, or cumulative probability (Kaplan-Meier-style) or as line plot with mean±SEM. Asterisks indicate statistical significance: *p < 0.05, **p < 0.01, ***p < 0.001. WT = wild type; KO = knockout. Animals were excluded for not meeting the required criteria to progress to the next phase.

In the DAT cohort, a linear mixed-effects model with fixed effects of session type (ethanol vs. sucrose) and genotype (WT vs. KO) was compared to a model with included genotype*session type interaction using a likelihood ratio test, which was found significant (χ²(1) = 56.77, *p* < .001). Pairwise comparisons revealed that KO animals showed significantly less responding than WT animals in the sucrose (-68%, β = 84.7, *SE* = 10.30, *t*(36.7) = 8.24, *p* < .001), but not ethanol (β = –9.28, *SE* = 9.38, *t*(29.8) = –0.99, *p* = .33) sessions. Within-group comparisons revealed higher responding for WT in sucrose than ethanol [β = -85.05, SE = 7.89, t(96.8) = - 10.78, p < .001].

For the SERT cohort, the analysis also revealed a significant genotype x session type interaction (χ²(1) = 5.64, *p* < .05). Pairwise comparisons revealed a significant reduction in operant responses for sucrose in SERT KO compared to WT rats (-28%, β = 32.8, *SE* = 9.84, *t*(26.9) = 3.33, *p* < .01), whereas no genotype differences were found in the ethanol sessions (β = 2.50, *SE* = 9.84, *t*(26.9) = 0.25, *p* = .802). Within-group comparisons revealed that both SERT KO and WT rats responded significantly more for sucrose than for ethanol (SERT WT: +91%; β = 108.0, *SE* = 8.8, t(14) = 12.30, *p* < .001; SERT KO: +92%; β = 78.0, *SE* = 8.8, t(14) = 8.86, *p* < .001).

### 3.5. DAT KO rats show delayed acquisition of ethanol- and sucrose-taking behavior, while SERT KO rats show delays only for sucrose

Acquisition speed of the operant task was evaluated as the number of days required to reach criterion performance, defined as ≥25 completed reward trials across two consecutive sessions, using Kaplan–Meier survival analysis for each of the three operant taking phases (Fig. 2C).

In the DAT cohort, no significant genotype difference was observed during ethanol-taking 1 (χ²(1) = 1.9, p = 0.2), with only two WT rats reaching criterion. However, during the subsequent sucrose-taking phase, DAT KO rats (x̄ = 33.13 ± 19.52 days) required significantly more sessions to reach criterion than WT rats (x̄ = 2.57 ± 0.49 days; χ²(1) = 14.20, p < .001). This delay persisted in ethanol-taking 2, where DAT KO rats (x̄ = 34.50 ± 7.30 days) again reached criterion significantly later than WT rats (x̄ = 6.67 ± 5.22 days; χ²(1) = 8.50, p = .003).

In the SERT cohort, no significant genotype effect was observed during ethanol-taking 1 (χ²(1) = 0.1, p = .7), with only one WT rat reaching criterion. In contrast, SERT KO rats (x̄ = 32.00 ± 27.16 days) required significantly more days to reach criterion than WT rats (x̄ = 3.40 ± 3.28 days) during the sucrose-taking phase (χ²(1) = 11.00, p < .001). This difference was not observed in ethanol-taking 2 (SERT WT: x̄ = 20.90 ± 14.08 days; SERT KO: x̄ = 23.00 ± 21.23 days; χ²(1) = 1.10, p = .304).

### 3.6. DAT and SERT WT and KO rats show reduced performance following sucrose-to-ethanol transition

To determine whether reward-taking behavior persists after a transition from sucrose to ethanol, we compared performance between genotypes for the first five ethanol sessions (Fig. 2D).

For the DAT cohort, linear mixed-effects models showed a significant main effect of genotype, with DAT KO rats displaying significantly lower ethanol-taking responding compared to DAT WT rats (β = –16.25, *SE* = 5.61, *t*(9.67) = –2.90, *p* = .016).

For the SERT cohort, in contrast, the analysis revealed no significant main effect of genotype (β = 4.66, *SE* = 10.82, *t*(10.90) = 0.43, *p* = .675).

### 3.7. DAT WT control rats and SERT cohort increased their ethanol taking following sucrose exposure

To examine whether prior sucrose exposure influenced subsequent ethanol-taking by comparing the average number of completed trials during the last five sessions of ethanol-taking 1 and the last five sessions of ethanol-taking 2 (Supplementary Figure 1B).

In the DAT cohort, a linear mixed-effects model with fixed effects of session type (ethanol 1 vs. ethanol 2) and genotype (WT vs. KO), and random intercepts for animal ID, revealed a significant genotype × session type interaction (χ²(1) = 23.07, p < .001).

Pairwise comparisons showed that WT rats significantly increased ethanol-taking after sucrose exposure (+41%; t(129.3) = 9.38, p < .001), whereas DAT KO rats showed no change (t(131.3) = –1.40, p = .976). In ethanol-taking 2, DAT KO rats completed significantly fewer trials than WT controls (–77.24%; t(26.70) = –5.39, p < .001), while no difference was observed in ethanol-taking 1 (t(19.40) = 1.88, p = .453).

In the SERT cohort, the same analysis showed no significant genotype × session type interaction (χ²(1) = 0.05, p = .822) and no main effect of genotype (β = –8.75, SE = 4.13, t(13.46) = –2.12, p = .053). However, there was a significant main effect of session type, with higher ethanol-taking in ethanol-taking 2 compared to ethanol-taking 1 (+56%; β = 9.38, SE = 3.92, t(13.17) = 2.40, p = .032).

## 4. Discussion

This study examined how constitutive deletion of the DAT and SERT influences motivation for natural (sucrose) and drug (ethanol) rewards, using both homecage and operant paradigms. By directly comparing sucrose and ethanol reward processing in DAT and SERT KO rats, we identified dissociable roles for the dopaminergic and serotonergic systems in modulating reward-driven behavior. DAT KO rats showed reduced sucrose preference and intake in the homecage (Fig. 1B), diminished Pavlovian responding for sucrose (Fig. 2B), delayed sucrose-taking acquisition (Fig. 2C), and complete disengagement from ethanol-taking after sucrose substitution (Fig. 2D), reflecting impaired reward valuation and effort allocation. SERT KO rats showed modest reductions in sucrose preference and intake (Fig. 1B), along with decreased operant responding for sucrose (Fig. 2B) and delayed sucrose-taking acquisition (Fig. 2C). They also exhibited reduced ethanol preference in the homecage (Fig. 1C) but maintained intact operant performance for ethanol (Figs. 2B–D). These findings highlight distinct contributions of dopamine and serotonin transporters to reward-driven behavior under varying levels of effort and motivational demand.

### DAT sucrose and ethanol sensitivity

DAT KO rats showed a clear reduction in both sucrose- and ethanol-driven behaviors, consistent with the hypothesis that chronically elevated extracellular dopamine induces a persistent reward-like state, or ‘reward saturation’ effect, which may diminish the motivational salience of external rewards by making them feel less necessary or impactful. This might reflect a condition where the brain’s reward circuitry is already tonically activated, reducing the drive to seek out additional rewarding stimuli. In line with previous findings, DAT KO rats had a significant reduction in sucrose preference (-27%, -13%) and intake (-43% at 4% sucrose) in the sucrose two-bottle choice, in the Pavlovian responding (--68.03%), and required at least 30 more sessions to reach acquisition criterion in the sucrose-taking task (Cinque et al., 2018; Mallien et al., 2022). In the homecage free-access paradigm, they also displayed reduced ethanol preference (−16%), though their ethanol intake (g/kg) was similar to WT. This is possibly due to lower bodyweight, which suggests reduced voluntary interests rather than pharmacological insensitivity and raises the possibility that similar pharmacological effects were achieved at lower intake levels.

During initial ethanol-taking (phase 1), few WT rats reached criterion, and DAT KO showed similarly low performance, indicating poor ethanol learning across genotypes in this phase. However, following the sucrose-to-ethanol switch (Phase 2), unlike WT rats, DAT KO rats exhibited a near-complete behavioral disengagement: lever pressing dropped to zero, and no animal reached acquisition criterion. This cannot be attributed to a global learning deficit, as they acquired the sucrose task (albeit more slowly), but rather suggests that ethanol lacked sufficient motivational value to sustain effortful behavior in the absence of sucrose. By comparison, while WT rats also reduced responding after the transition, most continued to engage and reached the criterion. These findings extend prior studies showing reduced ethanol intake in DAT knockdown mice and modest reduction in binge drinking in partial DAT deletion in rats (Bahi and Dreyer, 2019; Kuiper et al., 2023). Although contributing factors such as hyperactivity, motor stereotypies—observed in other studies (Cinque et al., 2018; Mallien et al., 2022)—and delayed learning capabilities, as suggested by previous findings showing impaired cocaine self-administration acquisition in DAT KO rats (Thomsen et al., 2009), may have contributed to slower task acquisition, they are unlikely to fully explain the reduced reward engagement observed here. Notably, ethanol’s acute effects on extracellular dopamine levels appear intact in DAT KO mice DAT (Mathews et al., 2006), suggesting that DAT is not essential for ethanol-induced dopamine elevation. Whether this holds true in rats—and with chronic exposure to ethanol or sucrose—remains unknown. Future studies using microdialysis or biosensors in DAT KO rats are needed to determine whether these rewards still evoke a dopaminergic response, or if compensatory mechanisms differ across species and reward types.

### SERT sucrose and ethanol sensitivity

SERT KO rats showed a reduction in reward-driven behaviors, particularly for sucrose. In the homecage two-bottle choice, sucrose preference was unchanged at 2% but slightly reduced at 4% (-5%), while sucrose intake was significantly lower at 4% (-46%). In the operant chamber, Pavlovian responding was reduced (−27.9%) and sucrose-taking acquisition was delayed by approximately 30 sessions, indicating subtle motivational impairments. These results align with previous reports of unchanged or slightly reduced sucrose preference and similar sucrose concentrations in males, but reduced intake under repeated long-access operant conditions (Bearer et al., 2009; Diniz et al., 2021; Karel et al., 2020; Olivier et al., 2008; Perona et al., 2008; Sun et al., 2024). In ethanol paradigms, SERT KO rats displayed markedly reduced preference (−34.8%) and intake (−28.5%) in the homecage, consistent with prior findings in SERT KO mice and rats, as well as with reduced ethanol self-administration after selective serotonin reuptake inhibitor treatment (Bearer et al., 2009; Haraguchi et al., 1990; Hussain et al., 2022; Kelaï et al., 2003; Lamb and Daws, 2013; Maurel et al., 1999). However, in operant settings, SERT KO rats performed comparably to WT controls: they showed intact Pavlovian responding, acquired ethanol-taking (Phase 1) at a similar rate, and continued responding after the sucrose-to-ethanol transition (Phase 2). This suggests that reduced serotonin function weakens ethanol’s reinforcing value under passive conditions but does not impair operant responding when ethanol is predictably available. Taken together, these results point to a context-dependent role of serotonin in reward-driven behavior— particularly under conditions involving effort, or affective load. This supports the concept of anxiety-related anhedonia, where heightened negative affect reduces motivational engagement despite intact hedonic valuation. Notably, SERT KO animals can perform well— or even outperform WT—in other tasks such as touchscreen-based paradigms under certain cue-driven conditions (Guo et al., 2021), reinforcing the view that motivational impairments in SERT KO rats are not global, but shaped by task structure and emotional context.

### Integration of findings

The findings of DAT and SERT KO rats combined highlight complementary roles for dopamine and serotonin in reward behavior: dopamine primarily supports effortful reward pursuit and value attribution, while serotonin modulates context-dependent engagement and affective tone. Supporting this dissociation, prior studies show that cocaine retains reinforcing effects when either DAT or SERT is absent, but not when both are deleted—suggesting that full expression of reward requires coordinated input from both systems (Hall et al., 2004; Rocha, 2003; Sora et al., 2001). This interplay may be critical for maintaining motivation under complex or shifting reward conditions.

### Clinical Implications

A preference for sweet tastes and heightened sensitivity to alcohol’s rewarding effects have been consistently linked to an increased risk of developing AUD (Carroll et al., 2002; Han et al., 2020; Kampov-Polevoy et al., 2004, 2001, 1999; King et al., 2016, 2014, 2011; McGraw et al., 2024; Morean and Corbin, 2010; Parker et al., 2020; Weafer et al., 2014). In this context, the observed reduction in sucrose and ethanol preference in DAT KO rats suggests that reduced DAT function may dampen reward sensitivity and potentially lower vulnerability to AUD-like behaviors. These findings support the therapeutic relevance of DAT inhibition, as seen with pharmacological agents like bupropion, which has shown efficacy in reducing alcohol intake in individuals with AUD and alcohol-preferring rats and may offer clinical benefit for individuals with high reward sensitivity profiles (Nicholson et al., 2018; Söderpalm et al., 2025). Finally, in movement disorders such as Parkinson’s disease, that are characterized by degeneration of nigrostriatal dopaminergic neurons, the loss of striatal DAT binding is associated with reward sensitivity (Barber et al., 2023).

Although SERT KO rats showed largely preserved sucrose preference, they displayed reduced ethanol preference and subtle motivational deficits under effortful conditions, suggesting mild reward insensitivity. These results align with preclinical studies linking serotonergic dysfunction to reduced ethanol intake (Sari et al., 2011). Clinically, SSRIs like fluoxetine and citalopram have shown mixed effects on alcohol use with some studies reporting reduced craving and intake and others suggesting poorer outcomes or increased drinking after discontinuation (Charney et al., 2015; Naranjo et al., 1992; Suárez et al., 2020). These discrepancies highlight the need to stratify patients by factors such as anxiety or serotonergic function. Our findings support the potential of serotonergic interventions for specific AUD subtypes rather than a one-size-fits-all approach.

### Limitations

While this study provides novel insights into genotype- and reward-specific motivational deficits, several limitations should be acknowledged. First, the use of constitutive knockouts precludes distinction between developmental and adult-onset effects, as lifelong absence of DAT or SERT likely induces compensatory changes. Future work using inducible or region-specific models could clarify the timing and locus of these effects. Second, we tested only one concentration of sucrose (10%) and ethanol (15%) in the operant tasks, limiting conclusions about dose-dependent reward sensitivity. Third, our operant tasks used fixed-ratio schedules, which may not fully capture motivational dynamics. Progressive ratio or effort-based paradigms could better assess effort allocation and breakpoint differences across genotypes. Furthermore, the SERT cohort completed a relatively low number in the first ethanol taking phase, masking a potential reduction due to a floor effect. Although based on a small sample (n < 10), the Kaplan–Meier analysis revealed a strong effect that should nonetheless be interpreted with caution due to limited power and potential uncertainty in the estimates. Lastly, the study included only male rats, despite known sex differences in monoaminergic function and reward processing (Warthen et al., 2020). Including females in future research is essential.

## Conclusion

In conclusion, this study reveals distinct roles for DAT and SERT in modulating reward-driven behavior. DAT deletion led to broad impairments in reward valuation and effortful engagement, consistent with a reward saturation hypothesis, while SERT deletion was associated selective deficits in sucrose motivation, and reduced ethanol preference, particularly under passive conditions. These findings underscore the importance of dopaminergic signaling for sustaining reward-driven effort and of serotonergic function in affective modulation and context-dependent reward engagement. Our results highlight how transporter-specific alterations influence behavioral responses to natural and drug rewards, offering insights into mechanisms underlying individual differences in substance use vulnerability.

---

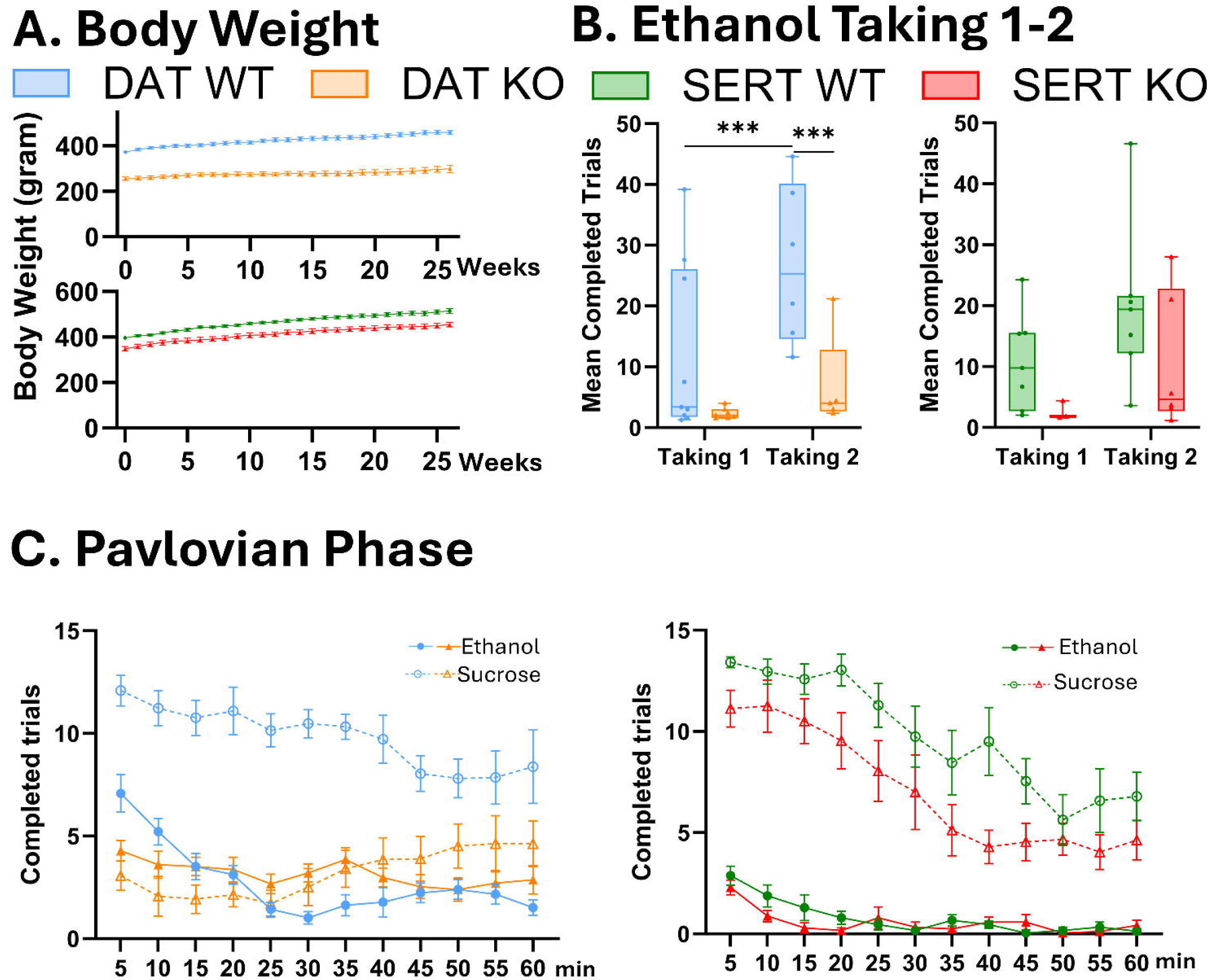

