## Supplementary Materials for "Different Sensitivity to Ethanol and Sucrose in DAT and SERT Knockout Rats"

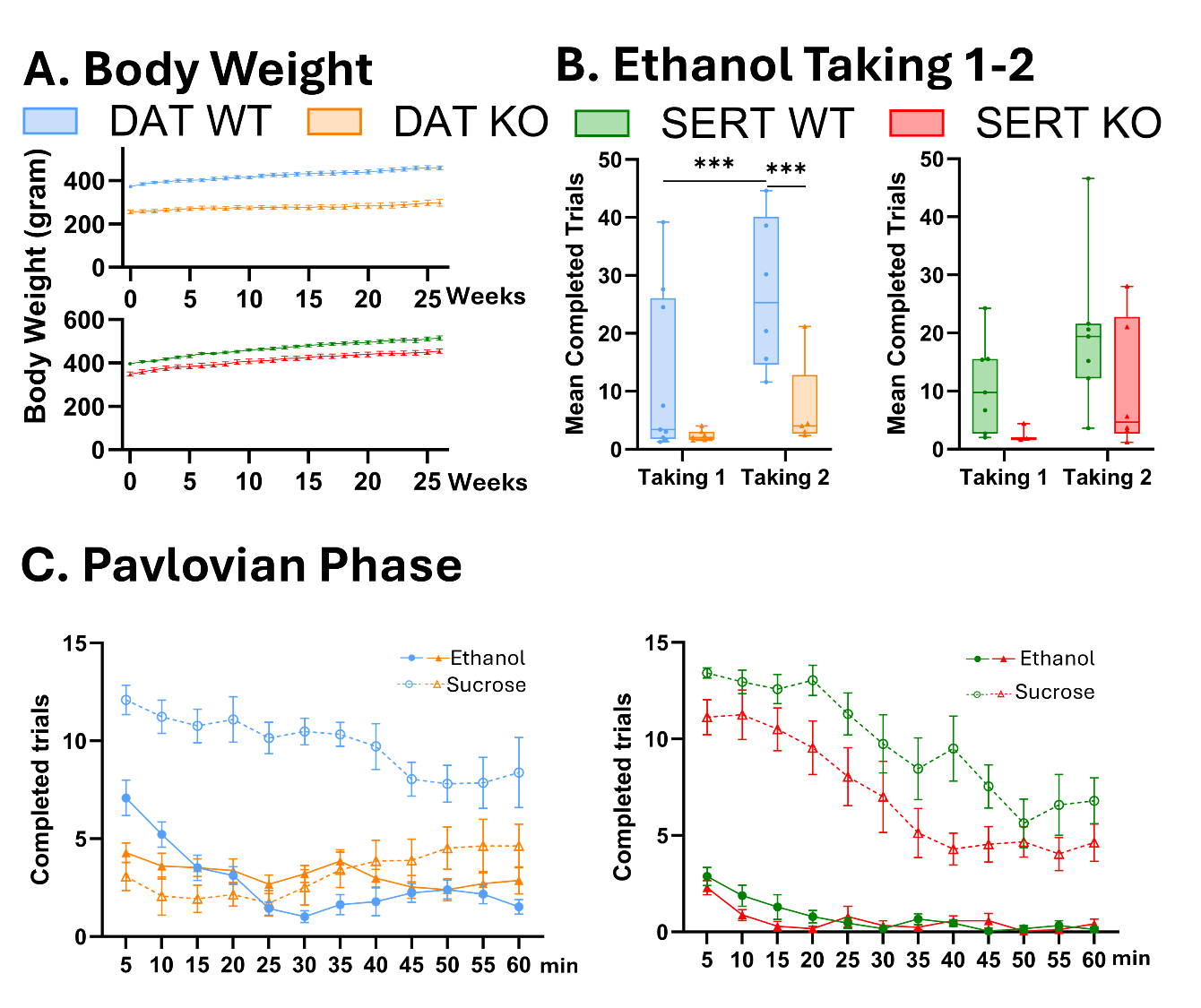

**Supplementary Figure 1. Bodyweight, and behavioral outcomes across DAT and SERT cohorts.** (A) Body weight curves across the 25-week study period. (B) Number of completed trials during ethanol taking 1 and 2 shown as the average number of completed trials per rat across the last 5 sessions of taking 1 and the last 5 sessions of taking 2. N = 7-10 DAT WT, 5-9 DAT KO, 7-8 SERT WT, 3-6 SERT KO. N = 10 DAT WT, 10 DAT KO, 8 SERT WT, 8 SERT KO. (C) Mean number of completed trials per 5-minute bin during the Pavlovian phase for sucrose and ethanol conditioning, averaged across the first three 1-hour sessions. N = 7-10 DAT WT, 9-10 DAT KO, 8 SERT WT, 8 SERT KO. Data are presented as boxplots (minimum to maximum) or line plot with mean±SEM. Statistically significant differences between groups are indicated by asterisks: *p < 0.05, **p < 0.01, ***p < 0.001. WT = wild type; KO = knockout.

**Supplementary Table 1.** Fixed effects measurements of linear model: bodyweight ~ group + (1|id) for all recorded bodyweight measurements (Sup. Fig. 1A).

| **DAT cohort** | | | | | |
| --- | --- | --- | --- | --- | --- |
| **Fixed Effect** | **Estimate** | **SE** | **df** | **t value** | **p-value** |
| Intercept | 386.49 | 8.636 | 20.33 | 44.75 | <.001 *** |
| genotypeKO | -145.98 | 11.847 | 18.01 | -12.32 | <.001 *** |
| **SERT cohort** | | | | | |
| **Fixed Effect** | **Estimate** | **SE** | **df** | **t value** | **p-value** |
| Intercept | 465.74 | 8.33 | 14.0 | 55.90 | < .001 *** |
| KO – WT | -52.86 | 11.78 | 14.0 | -4.49 | <.001 *** |

**Supplementary Table 2.** Fixed effects measurements of linear mixed model: preference ~ genotype + group + (1|id) of the preference (%) readout of the ethanol two-bottle choice.

| **DAT cohort** | | | | | |
| --- | --- | --- | --- | --- | --- |
| **Fixed Effect** | **Estimate** | **Std. Error** | **df** | **t value** | **p-value** |
| (Intercept) | 23.076 | 5.539 | 17.995 | 4.166 | <.001 *** |
| genotypeKO | -16.526 | 6.368 | 8.000 | -2.595 | 0.031868 * |
| session2 | 2.034 | 4.735 | 117.000 | 0.430 | 0.668257 |
| session3 | 6.012 | 4.735 | 117.000 | 1.271 | 0.206134 |
| session4 | 2.124 | 4.735 | 117.000 | 0.452 | 0.651781 |
| session5 | 6.602 | 4.735 | 117.000 | 1.394 | 0.165987 |
| session6 | 3.652 | 4.735 | 117.000 | 0.771 | 0.442001 |
| session7 | 2.241 | 4.735 | 117.000 | 0.474 | 0.636410 |
| session8 | 5.105 | 4.735 | 117.000 | 1.078 | 0.283245 |
| session9 | 9.451 | 4.735 | 117.000 | 1.995 | 0.049053 * |
| session10 | 19.001 | 4.735 | 117.000 | 4.013 | <.001 *** |
| session11 | 16.022 | 4.735 | 117.000 | 3.384 | 0.001171 ** |
| session12 | 22.457 | 4.735 | 117.000 | 4.743 | <.001 *** |
| session13 | 26.822 | 4.735 | 117.000 | 5.627 | <.001 *** |
| session14 | 12.577 | 4.735 | 117.000 | 2.657 | 0.008997 ** |
| **SERT cohort** | | | | | |
| **Fixed Effect** | **Estimate** | **Std. Error** | **df** | **t value** | **p-value** |
| (Intercept) | 23.904 | 4.318 | 31.909 | 5.536 | <.001 *** |
| genotoypeKO | -97.812 | 38.903 | 6.00 | -2.514 | 0.045640* |
| session2 | -28.995 | 48.841 | 91.00 | -0.594 | 0.554215 |
| session3 | 0.3493 | 48.841 | 91.00 | 0.072 | 0.943135 |
| session4 | -35.002 | 48.841 | 91.00 | -0.717 | 0.475429 |
| session5 | -19.888 | 48.841 | 91.00 | -0.407 | 0.684813 |
| session6 | -38.729 | 48.841 | 91.00 | -0.793 | 0.429860 |
| session7 | -26.209 | 48.841 | 91.00 | -0.537 | 0.592836 |
| session8 | 47.098 | 48.841 | 91.00 | 0.964 | 0.337443 |
| session9 | -18.628 | 48.841 | 91.00 | -0.381 | 0.703796 |
| session10 | 72.030 | 48.841 | 91.00 | 1.475 | 0.143719 |
| session11 | 114.496 | 48.841 | 91.00 | 2.344 | 0.021240* |
| session12 | 89.928 | 48.841 | 91.00 | 1.841 | 0.068845 |
| session13 | 246.229 | 48.841 | 91.00 | 5.041 | <.001 *** |
| session14 | 178.934 | 48.841 | 91.00 | 3.664 | <.001 *** |

**Supplementary Table 3.** Fixed effects measurements of linear mixed model: intake ~ genotype + group + (1|id) of the intake (g/kg) readout of the ethanol two-bottle choice.

| **DAT cohort** | | | | | |
| --- | --- | --- | --- | --- | --- |
| **Fixed Effect** | **Estimate** | **SE** | **df** | **t value** | **p-value** |
| (Intercept) | 1.9934 | 0.4845 | 14.485 | 4.115 | <.001 *** |
| genotypeKO | -0.3570 | 0.5894 | 8.000 | -0.606 | 0.561492 |
| session2 | 0.1434 | 0.3625 | 117.000 | 0.396 | 0.694348 |
| session3 | 0.3976 | 0.3625 | 117.000 | 1.097 | 0.274912 |
| session4 | 0.1923 | 0.3625 | 117.000 | 0.530 | 0.596784 |
| session5 | 0.3649 | 0.3625 | 117.000 | 1.007 | 0.316175 |
| session6 | 0.1643 | 0.3625 | 117.000 | 0.453 | 0.651929 |
| session7 | 0.2701 | 0.3625 | 117.000 | 0.745 | 0.457686 |
| session8 | 0.0010 | 0.3625 | 117.000 | 0.003 | 0.997699 |
| session9 | 1.0349 | 0.3625 | 117.000 | 2.855 | 0.005900 ** |
| session10 | 0.9055 | 0.3625 | 117.000 | 2.497 | 0.013967 * |
| session11 | 1.1469 | 0.3625 | 117.000 | 3.163 | 0.001987 ** |
| session12 | 1.3908 | 0.3625 | 117.000 | 3.835 | <.001 *** |
| session13 | 2.6259 | 0.3625 | 117.000 | 7.244 | <.001 *** |
| session14 | 1.3627 | 0.5093 | 117.000 | 2.675 | 0.008055 ** |
| **SERT cohort** | | | | | |
| **Fixed Effect** | **Estimate** | **SE** | **df** | **t value** | **p-value** |
| (Intercept) | 174.631 | 0.27228 | 14.05 | 6.414 | <.001 *** |
| genotypeKO | -0.50697 | 0.30979 | 6.00 | -1.637 | 0.15285 |
| session2 | -0.29540 | 0.23734 | 91.00 | -1.245 | 0.21647 |
| session3 | 0.03059 | 0.23734 | 91.00 | 0.129 | 0.89772 |
| session4 | -0.36993 | 0.23734 | 91.00 | -1.559 | 0.12255 |
| session5 | -0.32679 | 0.23734 | 91.00 | -1.377 | 0.17193 |
| session6 | -0.57836 | 0.23734 | 91.00 | -2.437 | 0.01676* |
| session7 | -0.34737 | 0.23734 | 91.00 | -1.464 | 0.14675 |
| session8 | -0.10170 | 0.23734 | 91.00 | -0.428 | 0.66931 |
| session9 | -0.28690 | 0.23734 | 91.00 | -1.209 | 0.22987 |
| session10 | 0.34535 | 0.23734 | 91.00 | 1.455 | 0.14908 |
| session11 | 0.62795 | 0.23734 | 91.00 | 2.646 | 0.00960** |
| session12 | 0.20508 | 0.23734 | 91.00 | 0.864 | 0.38983 |
| session13 | 0.72996 | 0.23734 | 91.00 | 3.076 | 0.00277** |
| session14 | 0.82614 | 0.23734 | 91.00 | 3.481 | <.001 *** |

**Supplementary Table 4.** Model comparison (full vs without interaction) using likelihood ratio test to assess the significance of the genotype * sessiontype (ethanol vs sucrose) interaction in pavlovian phase in operant chamber.

| **DAT cohort** | | | | | | | | |
| --- | --- | --- | --- | --- | --- | --- | --- | --- |
| **Model** | **No. Parameters** | **AIC** | **BIC** | **LogLik** | **Deviance** | **Chi²** | **df** | **p-value** |
| Completedtrials ~ genotype + sessiontype + (1\|id) | 5 | 1095.2 | 1108.6 | 542.60 | 1085.2 |  |  |  |
| Completedtrials ~ genotype * sessiontype + (1\|id) | 6 | 1040.4 | 1056.5 | 514.21 | 1028.4 | 56.773 | 1 | <.001  *** |
| **SERT cohort** | | | | | | | | |
| **Model** | **No. Parameters** | **AIC** | **BIC** | **LogLik** | **Deviance** | **Chi²** | **df** | **p-value** |
| Completedtrials ~ genotype + sessiontype + (1\|id) | 5 | 292.3 | 299.6 | -141.13 | 282.3 |  |  |  |
| Completedtrials ~ genotype * sessiontype + (1\|id) | 6 | 288.6 | 297.4 | -138.31 | 276.6 | 5.641 | 1 | 0.018* |

**Supplementary Table 5.** Estimated marginal means pairwise comparison (Bonferroni-corrected) for genotype*sessiontype interaction for linear mixed model of pavlovian phase in operant chamber.

| **DAT cohort** | | | | | |
| --- | --- | --- | --- | --- | --- |
| **Contrast** | **Mean**  **Difference** | **SE** | **df** | **95% CI**  **(Lower–Upper)** | **p-value** |
| WT – KO (Ethanol) | –9.2 | 9.28 | 29.8 | –28.9 to 10.5 | 0.3295 |
| WT – KO (Sucrose) | 84.7 | 10.30 | 36.7 | 63.9 to 105.5 | <.001*** |
| **SERT cohort** | | | | | |
| **Contrast** | **Mean**  **Difference** | **SE** | **df** | **95% CI**  **(Lower–Upper)** | **p-value** |
| WT – KO (Ethanol) | 2.5 | 9.84 | 26.9 | -17.7 to 22.7 | 0.8015 |
| WT – KO (Sucrose) | 32.8 | 9.84 | 26.9 | 12.6 to 53.0 | 0.0025** |

**Supplementary Table 6.** Fixed effects measurements of linear model: preference ~ genotype + session + (1|id) only for the 5 ethanol sessions following transition from sucrose taking (Fig. 2D).

| **DAT cohort** | | | | | |
| --- | --- | --- | --- | --- | --- |
| **Fixed Effect** | **Estimate** | **SE** | **df** | **t value** | **p-value** |
| (Intercept) | 39.546 | 6.050 | 40.982 | 6.536 | <.001*** |
| genotypeKO | -16.254 | 5.609 | 9.671 | -2.898 | 0.0164* |
| session2 | -0.023 | 0.969 | 41.010 | -0.147 | 0.884 |
| session3 | -1.310 | 6.969 | 41.010 | -0.188 | 0.8518 |
| session4 | -0.586 | 0.969 | 41.051 | -0.581 | 0.4000 |
| session5 | -11.530 | 7.139 | 41.475 | -1.615 | 0.1139 |
| **SERT cohort** | | | | | |
| **Fixed Effect** | **Estimate** | **SE** | **df** | **t value** | **p-value** |
| (Intercept) | 48.998 | 9.413 | 24.17 | 5.206 | <.001*** |
| genotypeKO | -4.328 | 11.117 | 10.65 | -0.389 | 0.70470 |
| session2 | -25.769 | 8.994 | 43.87 | -2.865 | 0.00637** |
| session3 | -28.448 | 9.476 | 44.76 | -3.002 | 0.00438** |
| session4 | -16.006 | 9.229 | 44.47 | -1.734 | 0.08978 |
| session5 | -22.340 | 9.229 | 44.47 | -2.421 | 0.01964* |

**Supplementary Table 7.** Model comparison (full vs without interaction) using likelihood ratio test to assess the significance of the genotype * sessiontype interaction of supplementary Fig. 1B comparison between ethanol taking 1 and ethanol taking 2 in operant chamber.

| **DAT cohort** | | | | | | | | |
| --- | --- | --- | --- | --- | --- | --- | --- | --- |
| **Model** | **No. Parameters** | **AIC** | **BIC** | **LogLik** | **Deviance** | **Chi²** | **df** | **p-value** |
| Completedtrials ~ genotype + sessiontype + (1\|id) | 5 | 1158.5 | 1173.4 | -574.28 | 1148.5 |  |  |  |
| Completedtrials ~ genotype * sessiontype + (1\|id) | 6 | 1137.5 | 1155.3 | -562.74 | 1125.5 | 23.066 | 1 | <.001*** |
| **SERT cohort** | | | | | | | | |
| **Model** | **No. Parameters** | **AIC** | **BIC** | **LogLik** | **Deviance** | **Chi²** | **df** | **p-value** |
| Completedtrials ~ genotype + sessiontype + (1\|id) | 5 | 192.21 | 198.31 | -91.107 | 182.21 |  |  |  |
| Completedtrials ~ genotype * sessiontype + (1\|id) | 6 | 194.16 | 201.47 | -91.081 | 182.16 | 0.0508 | 1 | 0.8217 |

**Supplementary Table 8.** Pairwise comparisons (Bonferroni-corrected) for genotype*sessiontype interaction of supplementary Fig. 1B comparison between ethanol taking 1 and ethanol taking 2 in operant chamber.

| **DAT cohort** | | | | | |
| --- | --- | --- | --- | --- | --- |
| **Contrast** | **Mean Diff.** | **SE** | **df** | **95% CI**  **(Lower–Upper)** | **p-value** |
| WT ethanol taking 1– KO ethanol taking 1 | 9.87 | 5.25 | 19.4 | –5.56 to 25.30 | 0.4526 |
| WT ethanol taking 1– WT ethanol taking 2 | –25.78 | 2.75 | 129.3 | –33.14 to –18.42 | <.001*** |
| WT ethanol taking 1– KO ethanol taking 2 | 5.51 | 5.61 | 23.9 | –10.62 to 21.64 | 1.0000 |
| KO ethanol taking 1– WT ethanol taking 2 | –35.65 | 5.46 | 22.1 | –51.47 to –19.83 | <.001*** |
| KO ethanol taking 1– KO ethanol taking 2 | –4.36 | 3.11 | 131.3 | –12.68 to 3.96 | 0.9764 |
| WT ethanol taking 2– KO ethanol taking 2 | 31.29 | 5.80 | 26.7 | 14.75 to 47.83 | <.001*** |
| **SERT cohort** | | | | | |
| No interaction | n.a. | n.a. | n.a. | n.a. | n.a. |
